# Local neuroplasticity in adult glaucomatous visual cortex

**DOI:** 10.1101/2022.07.04.498672

**Authors:** Joana Carvalho, Azzurra Invernizzi, Joana Martins, Remco J. Renken, Frans W. Cornelissen

**Author notes:** Correspondence: Joana Carvalho, Pre-clinical MRI laboratory, Champalimaud Centre for the Unknown, Avenida de Brasilia, 1400-038 Lisbon, Portugal. Contributed equally (Azzurra Invernizzi, Joana Martins).

## Abstract

The degree to which the adult human visual cortex retains the ability to functionally adapt to damage at the level of the eye remains ill-understood. Previous studies on cortical neuroplasticity primarily focused on the consequences of foveal visual field defects (VFD), yet these findings may not generalize to peripheral defects such as occur in glaucoma. Moreover, recent findings on neuroplasticity are often based on population receptive field (pRF) mapping, but interpreting these results is complicated in the absence of appropriate control conditions. Here, we used fMRI-based neural modeling to assess putative changes in pRFs associated with glaucomatous VFD. We compared the fMRI-signals and pRF estimates in participants with glaucoma to those of controls with case-matched simulated VFD. We found that the amplitude of the fMRI-signal is reduced in glaucoma compared to control participants and correlated with disease severity. Furthermore, while coarse retinotopic structure is maintained in all participants with glaucoma, we observed local pRF shifts and enlargements in early visual areas, relative to control participants. These differences imply that the adult brain retains local neuroplasticity. This finding has translational relevance, as it is consistent with VFD masking, which prevents glaucoma patients from noticing their VFD and seeking timely treatment.

## 1. Introduction

Damage to the visual system, for instance due to a retinal or a cortical lesion, or a development disorder, results in visual field (VF) loss (scotomas) and deprives the visual cortex from its normal input ^1–5^. When such damage occurs early in life, the visual cortex has the capacity to modify its retinotopic organization to compensate for vision loss: neurons affected by the lesion become responsive to other parts of the VF ^1, 5–10^. Whether or not the adult human visual system retains this plasticity is a deeply debated issue. Some studies suggest that the adult visual cortex can compensate for visual damage by re-scaling and displacing receptive fields (RFs) towards spared regions of the visual field ^11–14^. This would result in partially restoring the visual input. Other studies found a remarkable degree of stability of the primary and extrastriate areas of the adult visual cortex following central retinal lesions acquired later in life ^15, 16^.

Thus far, the vast majority of studies on cortical reorganization have addressed disorders that affect foveal vision, i.e. the central part of the VF ^12, 13, 15, 17, 18^. However, the findings of these studies are controversial; some studies support plasticity of the visual system ^11–14^ while others favor its stability ^15, 16^. Furthermore these findings may not apply throughout the VF, such as its periphery, as the neuronal density devoted to process foveal inputs is exponentially larger than the one allocated to peripheral information. Still, it is important to also understand cortical plasticity in peripheral parts of the VF. In the ophthalmic disease glaucoma, VF loss typically starts in the periphery. This slowly progressing neurodegenerative disease is the second leading cause of permanent visual impairment among the elderly worldwide ^19^. Detection of the impairment by the patient themselves is often delayed due to the perceptual masking of the VFD by their own brain, also referred to as “filling-in”^20, 21^. Although this phenomenon is also observed in normal perception, in patients it is postulated to be a byproduct of neuroplasticity ^22^. Consequently, the mechanisms underlying functional reorganization in glaucoma might differ from those involved in diseases that affect central vision. Thus far, only few studies investigated functional cortical changes associated with glaucoma ^23–25^. Zhou and colleagues investigated retinotopic visual function in the visual cortex of POAG participants using wide-view visual presentation (up to 55 degrees). They found an enlarged representation of the parafovea and a larger cortical magnification of the central visual representation in the visual cortex of POAG participants compared to control participants. They interpreted these changes as evidence of cortical remapping ^25^. However, whether the mere presence of differences in size and position of RFs is evidence of cortical reorganization has been questioned, as similar changes occur with simulated (artificial) scotomas ^15, 17, 18, 26–28^. Consequently, it remains unresolved whether such changes in representation are the result of reorganization or simply the consequence of the damage at the level of the eye changing the input that reaches the cortex. Therefore, to fully address the issue of retained adult plasticity in glaucoma, it is essential to include studies on participants with peripheral VFDs, to use high-precision analyses, and to include control conditions that account for the visual deprivation due to natural scotomas.

For these reasons, in this study, we investigated how glaucoma affects the functional organization of the visual cortex using fMRI in combination with advanced neural modeling. We assess the fMRI responses of participants with glaucoma vis-a-vis those of age-matched control participants with a matched, simulated VFD. This is crucial, as it will allow us to rule out that any observed changes in the pRFs are merely the result of the altered visual input ^15, 17, 25, 29, 30^. Therefore, for each participant with glaucoma, a matched control participant observed the visual stimuli with a simulated scotoma (SS) designed to mimic the glaucoma participant’s reduced visual sensitivity, as assessed using standard automated perimetry (SAP). Moreover, we assessed retinal thickness using Optical Coherence Tomography (OCT) and applied fMRI-based VF mapping techniques, based on both standard pRF mapping and an advanced variant called micro-probing ^31^.

To preview our results, while the coarse retinotopic reorganization of the visual cortex was maintained in the participants with glaucoma, we found a reduction in its BOLD responsiveness and differences in the distributions of estimated pRF sizes and positions compared to those in control participants with a matched SS. In other words, the observed pattern of reorganization of the visual cortex in glaucoma appears not to be a mere consequence of the reduced visual input due to the retinal scotoma. Moreover, when expressed in terms of the changes in pRF size and location, we find that the degree of reorganization correlates with the severity of the glaucomatous VF damage. In addition, fMRI-based VF reconstructions showed that glaucoma participants exhibit a lower VF sensitivity compared with controls with SS and local differences between fMRI-based VF sensitivity patterns from those obtained via SAP. Together, our findings support that the adult visual cortex retains a spatially localized capacity to functionally reorganize.

## 2. Results

### 2.1 Glaucoma affects the BOLD modulation in the visual cortex

Figure 1A shows the BOLD modulation as a function of eccentricity for participants with glaucoma and for the control participants with and without a simulated scotoma, controls SS and controls NS, respectively. At all eccentricities, the BOLD modulation is reduced in the participants with glaucoma compared to both control conditions (F(2,31)=12.98, p<0.0001, FDR-corrected). There was no evidence for differences in the BOLD modulation of individual visual areas (F(2, 31)=2.67, p=0.07) nor for an overall effect of eccentricity (F(12, 341)=0.49, p=0.92). The slightly higher foveal responsiveness of the control participants in the SS compared to the NS condition caused a significant interaction of group and eccentricity (F(12, 341)=2.05, p=0.02).

**Figure 1.**
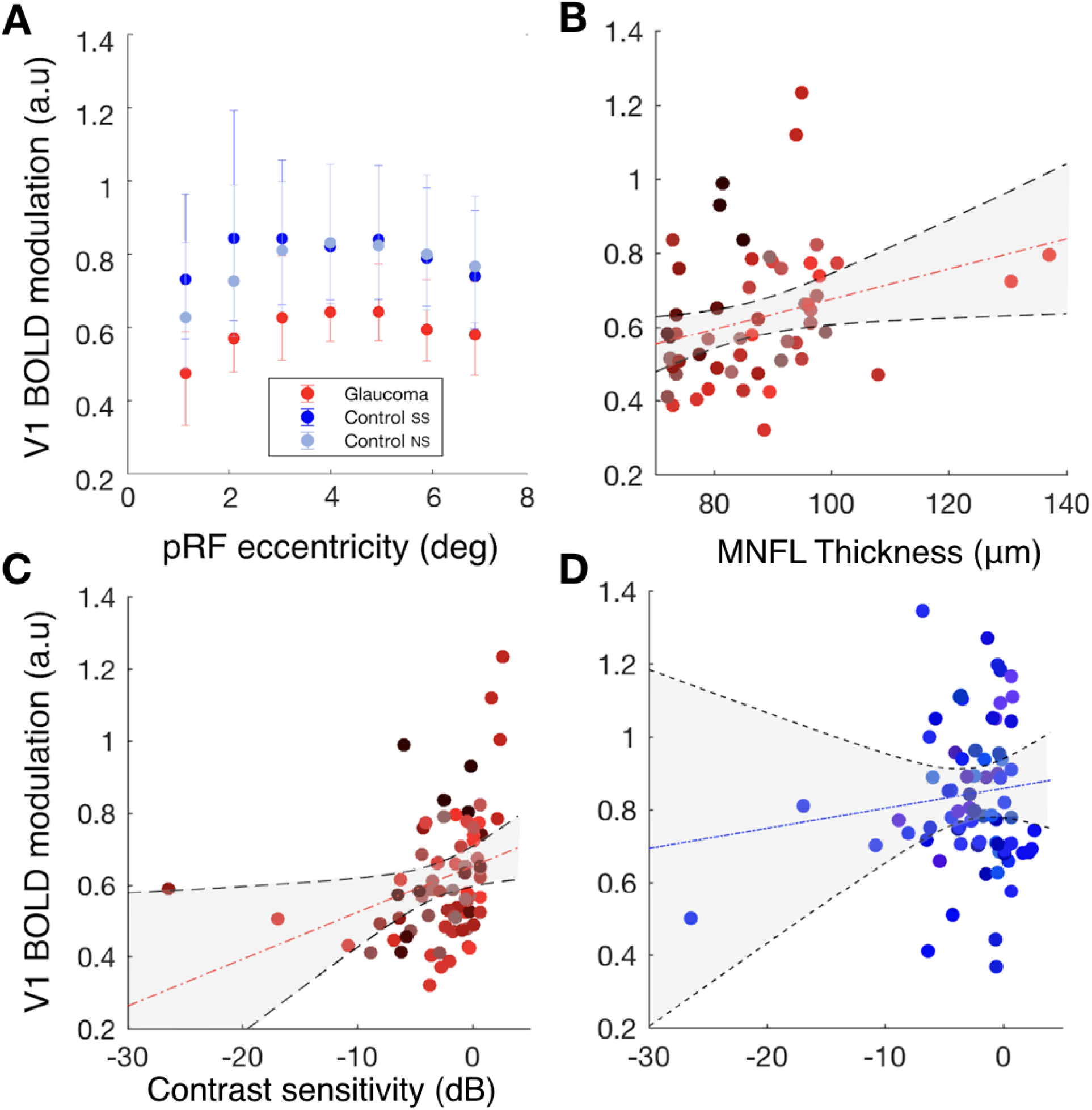
- V1 BOLD modulation varies between participants with glaucoma and control participants and correlates with disease severity. A: V1 BOLD modulation, defined as a function of eccentricity for glaucoma (red) and control participants with (dark blue) and without a simulated scotoma (light blue). The BOLD modulation was binned in 1 degree bins. The bars represent the 95% confidence interval. B: Correlation of the BOLD modulation of participants with glaucoma obtained from individual quadrants with MNFL thickness (calculated by averaging the values of the macula of both eyes). Each data point is from a separate quadrant of an individual participant with glaucoma. C and D: Correlation of the BOLD modulation from separate quadrants with the mean deviation (MD) of both eyes combined (the max between the MD of the two eyes) for glaucoma and controls, respectively. Each data point is from an individual VF quadrant.

Figures 1B and 1C show the average BOLD modulation as a function of contrast sensitivity and macular nerve fiber layer (MNFL) thickness, respectively. Data are plotted per individual VF quadrant. For both parameters, we find significant correlations with average BOLD modulation (contrast sensitivity: r^2^ = 0.31, p = 0.006; MNFL thickness r^2^ = 0.31 p = 0.007). As shown in Figure S1, the MNFL thickness correlation is also present for visual areas V2 (r^2^ = 0.34 p = 0.0024) and V3 (r^2^ = 0.35 p = 0.0017) while a significant correlation of BOLD vs contrast sensitivity is present for V3 but not for V2 (V2: r^2^ = 0.2, p = 0.08; V3: r^2^ = 0.22, p = 0.05). Figure 1D shows the V1 BOLD modulation of control participants in the SS condition as a function of simulated contrast sensitivity. The correlation is not significant (r^2^ = 0.08, p = 0.44). As shown in Figure S1, this was neither the case for V2 (r^2^ = −0.01, p = 0.92) nor for V3 (r^2^ = 0.09, p = 0.42).

### 2.2 Large scale organization of the visual cortex is preserved in glaucoma

Figure 2A shows the pRF properties (eccentricity, polar angle and size) projected on the inflated brain mesh, obtained for participant pair P03 consisting of a participant with glaucoma (G03) and matched control participant (C03). The latter performed the experiments both without (NS) and with (SS) a simulated scotoma matched to the scotoma of G03. Despite the severe glaucomatous VFD, represented by the VF plots measured using SAP in panel 2B, the global retinotopic organization of the visual cortex is preserved in G03. Figure S2 shows the retinotopic maps of additional glaucoma participants with different VFDs. Panels 2D and 2E show that the eccentricity and polar angle histograms exhibit the same pattern for G03 and C03 in both the NS and SS conditions, and only local differences can be observed. Regarding pRF size, overall we found no significant differences between G03 and C03 SS and NS. Still, as shown in Panel 2F, at more peripheral locations (5° to 7°) which are more affected by the VFD both G03 and C03 (SS condition) show larger pRFs sizes than C03 (NS condition). As expected, G03 and C03 (SS and NS) show an increase of pRF size with eccentricity. Panel 2C shows various single-voxel time series for the glaucoma and control participant. The voxel locations are indicated by the colored arrows in panel 2A. The top graph of panel 2C shows the time series recorded in two voxels. One voxel is located in the SPZ of G03 (red arrow) which, in this case, is situated in the periphery of the lower left quarter field. Of relevance, according to SAP, G03 still had residual sensitivity within their SPZ. The second time series was recorded in a voxel with a (mirrored) pRF position in the upper left visual quarter field. Therefore, this was located in the contralateral hemisphere (orange arrow). Not so surprising, the BOLD modulation of the voxel inside the SPZ (dark red line) is substantially lower than the one located in the contralateral hemisphere (orange line). Moreover, this decrease was substantially larger than expected based on the reduced stimulation that is the consequence of the VFD, as mimicked by the simulation in C03 (SS), as shown in the bottom graph of panel 2C which depicts the time series of a voxel located in the simulated SPZ of C03 in SS and NS conditions (dark and light blue lines and arrows, respectively). These examples illustrate that, while there may be a decrease in BOLD responsiveness within the SPZ in glaucoma, the large-scale organization of the visual cortex can still be preserved. Further examples of individual data can be found in Figure 3, panels A-D and Figure S2.

**Figure 2.**
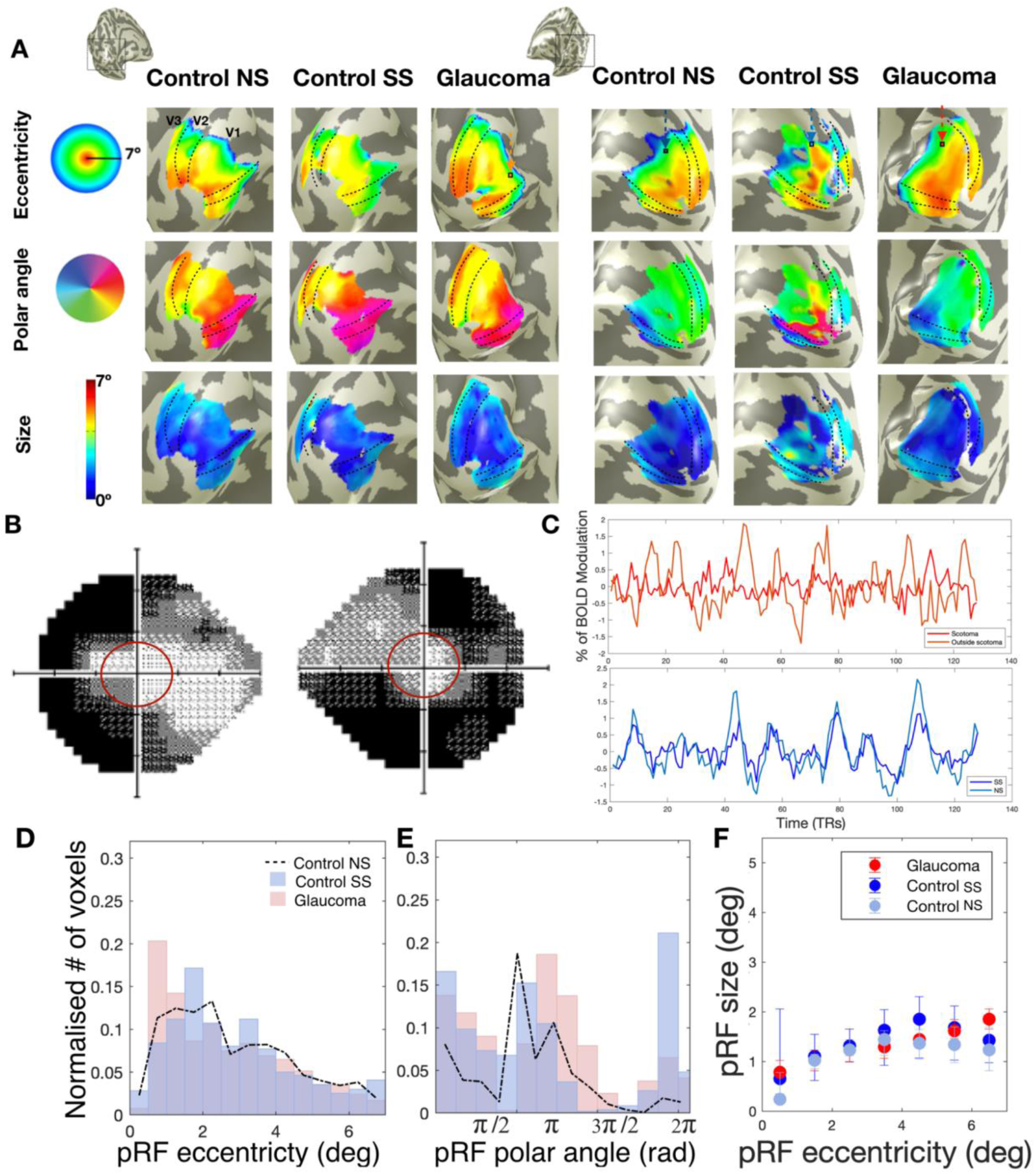
Preserved cortical organization in glaucoma. A: Eccentricity, polar angle, and pRF size maps obtained for a participant with glaucoma (G03), and their respective control participant (C03) with (SS) and without (NS) a matched simulated scotoma. The maps were obtained using an explained variance threshold of 0.1 The dashed lines delineate the visual areas V1, V2 and V3. B: VFs for the left and right eye of the participant with glaucoma. The red line corresponds to the VF that can be mapped using fMRI (7 degrees radius), C: Example time series (arrows in the top row of figures of panel A show the voxel locations). The upper panel in C shows the time series of a voxel of G03 in their SPZ (red arrow) and of a voxel with a similar pRF position but located in the contralateral hemisphere (orange arrow). The lower panel in C shows for C03 the time series of a voxel located in their simulated SPZ during the NS (dark blue arrow) and SS conditions (light blue arrow), respectively. D and E: Histogram of the normalized number of responsive voxels (i.e.VE>15%), as a function of eccentricity and polar angle, respectively. G03 is depicted in red and C03 in the simulation condition in blue (Control SS, i.e. the simulation matched to G03). C03’s results in the no simulation (NS) condition are depicted by the dashed black line. E: PRF size as a function of eccentricity for G03 (red) and C03 in the conditions NS (light blue) and SS (dark blue). The error bars correspond to the 95% confidence interval.

**Figure 3.**
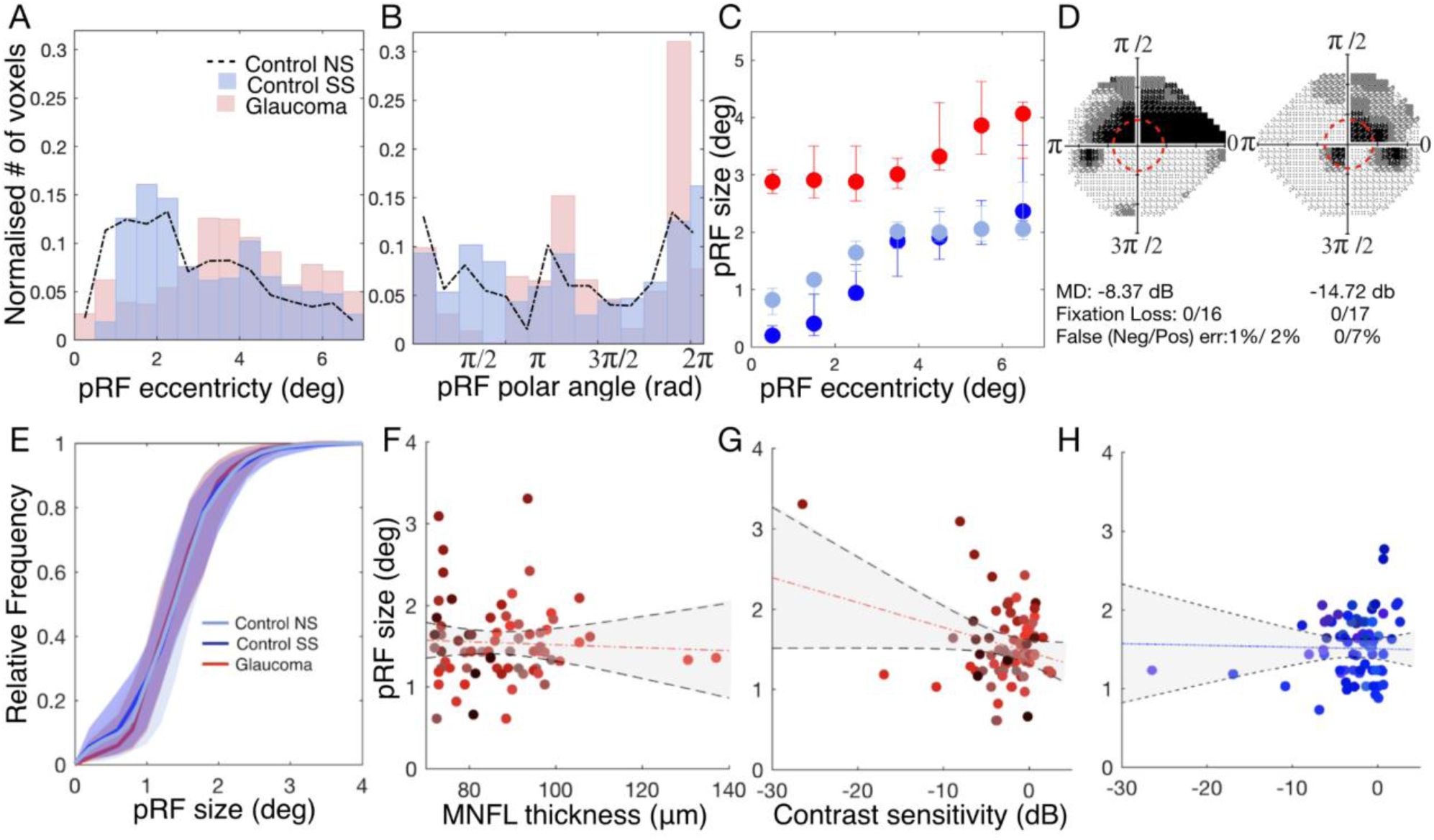
V1 pRF size for participants with glaucoma and control participants. Top row: Example of single participant data: V1 pRF properties differ between participants G01 and C01. A and B: Histogram of the normalized number of responsive voxels, those whose VE>15%, as a function of eccentricity and polar angle, respectively. The glaucoma participant (G01) is depicted in red and the control participant (C01) in the simulation condition in blue (SS; simulation matched to G01). The control participant’s results in the no simulation (NS) condition are depicted by the dashed black line. C: VFs for the left and right eye. D: Cumulative distribution of the V1 pRF sizes for the glaucoma participant (red) and the control participant in the conditions NS (light blue) and SS (dark blue). E: PRF size as a function of eccentricity for the participant with glaucoma (red) and the control participant in the conditions NS (light blue) and SS (dark blue). The error bars correspond to the 95% confidence interval. Bottom row: Group level analysis. E: Mean cumulative distributions of the pRF size in V1 for the glaucoma participants (red) and the control participants (NS: light blue, SS: dark blue). F and G: PRF size obtained for the glaucoma participants as a function of the contrast sensitivity and RNFL of the macula, respectively. Data are shown per VF quadrant. Different colors denote the different participants. The dashed red line depicts the linear fit while the black dotted lines and shaded areas correspond to the 95% CIs. Panel H is a similar plot as panel G but for the control participants (SS condition).

### 2.3 Local pRF properties differ between participants with glaucoma and control participants with simulated scotoma

#### 2.3.1 Regions of lower contrast sensitivity are associated with larger pRFs

Next, we investigated if, at a finer-scale, the organization of the visual cortex in participants with glaucoma. The visual field defects (VFD) in our cohort of participants with glaucoma differed substantially. This makes it less meaningful to simply average their data, when investigating changes in pRF properties.

The top panels of Figure 3 show an analogous analysis to the one shown in Figure 2, but for the participant pair P01, consisting of glaucoma participant G01 and their associated control participant C01. The projections of the pRF properties in the inflated brain mesh for G01 can be found in Figure S2. Panels 3A and 3B indicate that while for C01, the pRF distributions in the NS and SS conditions are similar, for G01 the normalized number of pRFs located in the scotomatic region (the binocular scotoma of G01 is primarily located in the upper left quadrant defined by a polar angle between 0 and 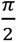, Panel 3D) is reduced compared to that in the SS condition of C01. Figure S3 shows the normalized distributions for the remaining participant scotoma-control pairs. Furthermore, Figure 3C shows that G01 exhibits larger pRFs than C01 which are not restricted to the scotomatic area but are present throughout the entire (central) VF. For completeness, we present the analysis of the variation of the pRF size as function of the polar angle in figure S4, which also shows that the glaucoma participant has larger pRFs sizes across the entire visual field when compared to the control participant, but with and without the simulation (SS and NS). However, such larger pRFs were not consistently found in all glaucoma participants. For example, in G03 depicted in Figure 2, the pRF size maps provide little evidence for deviations in pRF size. In fact, as we will report further on, heterogeneity rules. We compared the normalized distribution of the pRF estimates between each glaucoma participant and their respective control, while we found deviations in all glaucoma participants, in some this primarily affected pRF size, while in others this was mainly reflected in the pRF location, Figures S7, S8 and S9. In particular figure S7 shows that glaucoma heterogeneously affects pRF size, evident from multiple observable patterns. The majority of glaucoma participants (10 out of 19) show larger pRFs compared to the matched controls (evident from a negative deviation for smaller pRFs and a positive deviation for larger pRFs, e.g P01). Others show the opposite pattern and have smaller pRF sizes compared to the matched controls (evident from a positive deviation for smaller pRFs and a negative deviation for the larger ones, e.g. P19). Yet, others show no significant differences at all (e.g. P15). However when grouped together, on average, there is no significant difference between the V1 pRF size of participants with glaucoma and the control participants with and without SS (t(60))=1.63, p=0.10), Figure 3E. Figure S3 shows that also for visual areas V2 and V3 there were no such differences (t(60))=-0.65, p=0.52 and t(60))=-0.26, p=0.80, respectively).

In order to perform group analysis taking into account the heterogeneity of the VFDs of glaucoma participants, we correlated the average pRF size per quarter field with the metrics obtained via ophthalmic tests (contrast sensitivity measured with SAP and MNFL thickness measured with OCT). Panel 3F shows that for the participants with glaucoma, the average V1 pRF size per quarterfield did not correlate with MNFL thickness (r^2^=0.008, p=0.7) while panel 3G shows that the average V1 pRF size per quarterfield did correlate with SAP VF sensitivity (r^2^=-0.28, p=0.02), with larger pRFs in case of lower contrast sensitivity. Figure 3H shows that in controls with simulated visual field defects, this effect was not found (r^2^=-0.02 p=0.84). Similar patterns are present for visual areas V2 and V3 (Figure S4). These results suggest enlargement of pRFs as a possible mechanism that compensates for the effect of a VFD, which would be in line with the finding of increased spatial integration as found with psychophysics ^32, 33^.

#### 2.3.2 Variation in average pRF location between glaucoma participants and controls SS correlated associated with contrast sensitivity loss

Figures S8 and S9 show the deviations in pRF eccentricity and polar angle between participants with glaucoma and the respective control participants. In all 19 glaucoma-control (SS) participant pairs, the distributions differed significantly from baseline in at least one of the bins. However, the location of these differences was highly variable between participants. Consequently, when evaluated at the group level, these differences tend to cancel each other out. This is confirmed in figures 4A and 4B, which show that when aggregated across all the participants with glaucoma and the control participants (SS), there are hardly any differences in the average normalized histograms of pRF eccentricity and polar angle.

**Figure 4:**
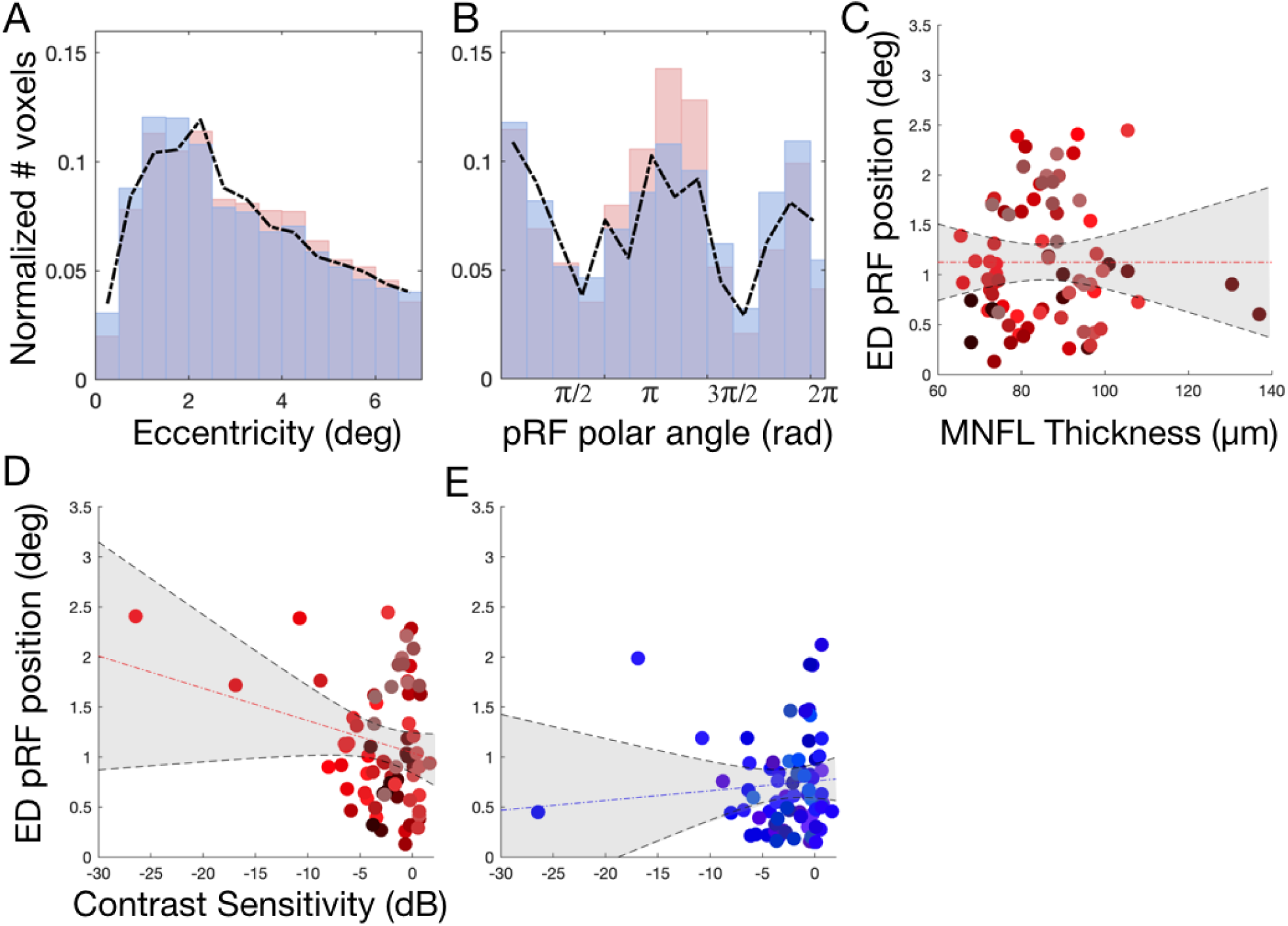
V1 pRF properties differ between participants with glaucoma and control participants. Panels A and B: Histogram of the normalized number of responsive voxels, those whose VE>15%, averaged across all the participants, as a function of eccentricity and polar angle, respectively. The average of all the glaucoma participants is depicted in red and the average of control participants SS in blue. The dashed black line depicts the histogram of the control participant’s in the no simulation (NS) condition. C and D: Correlation between the Euclidean Distance (ED) between the average pRF position per VF quadrant obtained for Glaucoma and Controls SS and the contrast sensitivity and MNFL, respectively. E: Analogous correlation to panel D but here the ED is calculated between Controls SS and Controls NS. Data are plotted per VF quadrant. Different colors denote the different participants. The dashed red line depicts the linear fit and the black line and shaded area corresponds to the 95% CI.

Importantly, at the individual level, we show that Euclidean Distance (ED) between the average position calculated per quarter field for glaucoma and respective control SS participants correlated significantly with contrast sensitivity (r^2^=-0.23; p=0.049, panel D) but not with macular thickness (r^2^=0.08; p=0.48, panel C). The within-participant ED between the SS and NS conditions does not correlate significantly with the simulated loss of contrast sensitivity (r^2^=-0.00003; p=0.99, panel E). Furthermore, this analysis is corroborated with a more complex, detailed and locally specific analysis, based on local deviations of the pRF properties distributions between participants with glaucoma and control participants SS. In line with Figure 4, Figure S10 shows that the pRF position deviations between participants with glaucoma and control participants SS correlated significantly with contrast sensitivity (r^2^=-0.24; p=0.004, panel A) but not with macular thickness (r^2^=-0.04; p=0.74, panel C). The deviation between control SS and NS does not correlate significantly with the simulated loss of contrast sensitivity (r^2^=-0.1; p=0.39, panel B). Figure S3 shows that the pRF parameter distributions are consistent across participants.

### 2.4 Glaucoma affects the visual system beyond the eye

To investigate whether: a) the brain representation of the VF matches the one measured at the level of the eye and b) there is a higher sampling of the SPZ that could explain some perceptual phenomena characteristic of glaucoma, such as predictive masking, we compared the reconstructed VF maps of the participants with glaucoma with the control SS participants. Figure 5 shows an example of the reconstructed VF of a pair of participants (glaucoma and the respective control SS and NS) and the SAP outcome of the right and left eye of the participant with glaucoma. According to SAP, the upper right quadrant is functionally blind (contrast sensitivity <-32 dB). For the glaucoma participant, the VF reconstruction nicely overlaps with the SAP tests, showing a reduced VF sensitivity to the upper right quadrant. Nevertheless, this reduction is not as strong as expected based on SAP, in particular in the periphery. In addition, although to a lesser extent, also the control SS participant shows a reduction in VF sensitivity in the quadrants most affected by the SS. Importantly, such reduction in VF sensitivity is not present in the control participant (NS). Furthermore, the VF reconstruction results are also in line with the pRF analysis shown in Figure 2. This confirms that: 1) the VF reconstruction techniques are accurate and allow to retrieve VF sensitivity in glaucoma; and 2) that the SS have the desired effect although at a smaller scale. Importantly, the real and SS scotomas become smaller with visual hierarchy which can explain the predictive masking of the scotomas experienced by glaucoma patients.

**Figure 5.**
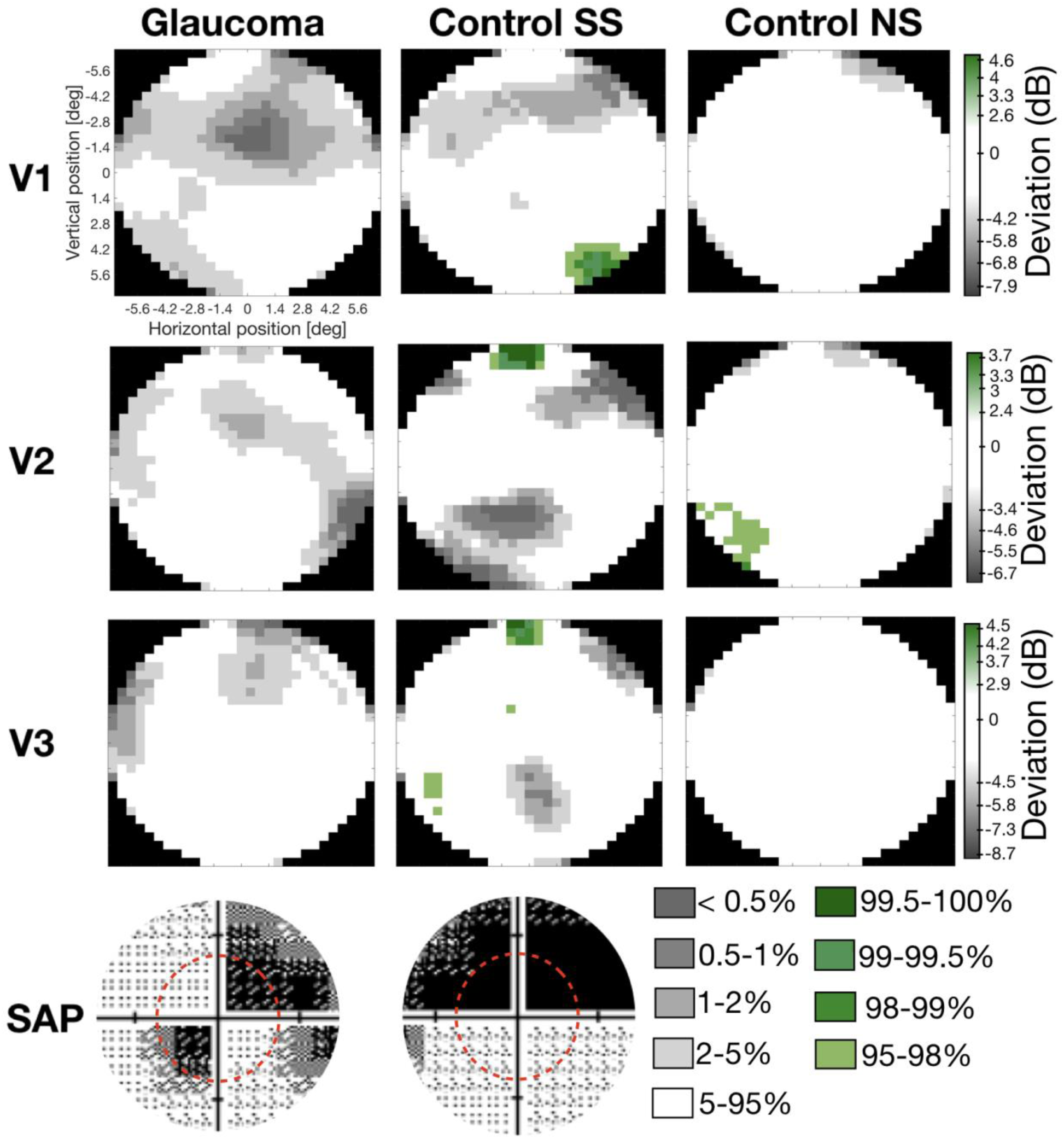
Comparison of the visual field reconstructions for visual areas V1, V2 and V3 of a participant glaucoma (G10) and the matched control participant (C02) with (SS) and without simulation (NS). The visual field reconstructions are based on the deviation between Glaucoma and Control SS from the average controls NS group (excluding the control participant with the SS matched to the glaucoma patient). The color code corresponds to the deviation to the CI, in particular regions of the VF within 5-95% CI are coded in white while the gray and green tones correspond to the < 5% and >5% normal limits, respectively. The bottom panel shows the glaucoma participant SAP test and dashed red line corresponds to the VF that could be mapped using fMRI.

When examining the VF reconstructions (Figure 6) across the pairs of the participants with glaucoma and their controls with matched simulation (SS) it is clear that the VF sensitivity in glaucoma participants is overall lower than in controls SS. The majority of the controls SS do not show a marked scotoma. This related with the fact that: a) only one glaucoma participant has a binocular scotoma with a sensitivity below -15dB (which is the threshold to be considered blind that portion of the VF) within the FOV of stimulation within the scanner; and b) a reduction in contrast does not affect significantly the pRF estimates ^34^. The differences between the glaucoma and control SS participants, suggests that either SS based on contrast sensitivity measured by SAP cannot accurately reproduce the real scotomas, or that the damage caused by glaucoma goes beyond the eye and affects the brain. Second different patterns are observed, while some glaucoma participants exhibit a lower VF sensitivity mapped via fMRI compared to what was measured with SAP, i.e.: P2, P5, P6, P7, P16, P17, for others the reverse pattern in observed, i.e P1, P3, P4, P8, P11, P19. This suggests that glaucoma participants develop different strategies to cope with their VFD and that the effect of glaucoma goes beyond the eye. Some controls SS show increased the VF sensitivity compared to the baseline controls NS (green regions in the VF reconstruction graphs), this increased VF sensitivity may reflect first the variation in fMRI based VF reconstructions and the effect of SS inducing short-term pRFs shifts directed to perceptually mask the scotoma ^35^.

**Figure 6.**
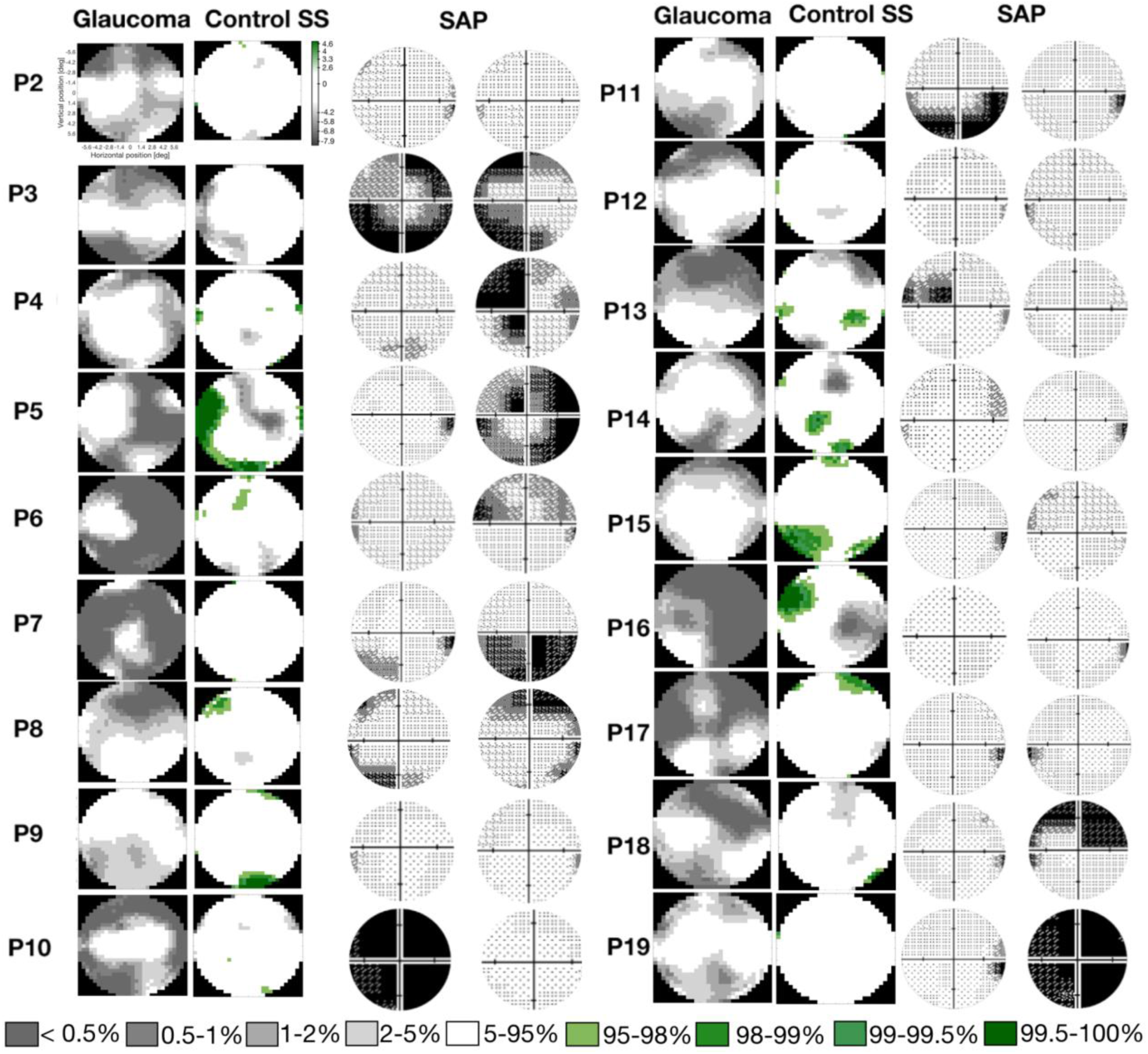
Comparison of the V1 visual field reconstruction between glaucoma and control SS participants. VF reconstruction glaucoma-control participants SS pairs and the SAP tests for the left and right eye are shown. Note that to compare the binocular reconstructed VF with the monocular HFA outcomes, the highest contrast sensitivity from both eyes should be considered.

## 3. Discussion

In this study, we used fMRI in combination with model-driven analyses to quantify changes in cortical functional organization in participants with the ophthalmic disease glaucoma. Various of our findings suggest that visual cortex functioning is altered in glaucoma, which could be indicative for cortical plasticity. Our main findings were: 1) cortical areas retained their coarse retinotopic organization, consistent with what has been described previously ^15, 36^; 2) marked differences in the magnitude of the BOLD response that cannot simply be explained by reduced contrast sensitivity; 3) local differences in the size and position of pRFs; and 4) notable differences in the fMRI-based VF reconstructions obtained for participants with glaucoma compared to those for matched control participants with similar but simulated visual loss. Our findings suggest that glaucoma is associated with limited and local functional remapping in the early visual cortex. These local neural reconfiguration patterns are highly variable between participants with glaucoma and likely depend on the preserved VF. Such local changes may be interpreted as attempts by the visual system to compensate for loss that results from the neural damage. Moreover, the observed pRF changes are consistent with predictive masking of the scotomas and may have implications for future treatments. Below, we discuss our findings and interpretation in detail.

In participants with glaucoma, throughout their entire VF, the magnitude of the BOLD modulation is reduced when compared to control participants with matched simulated VFDs. The comparison to the latter indicates that these changes cannot be explained by reductions in contrast sensitivity reducing the strength of the signals reaching the cortex. In particular, we found a reduction in the BOLD modulation at larger eccentricities (>5 deg), consistent with the notion that glaucoma primarily affects peripheral vision. Nevertheless, also at eccentricities <5 deg the modulation was decreased. Moreover, when analyzed per quadrant of the VF, the modulation of the fMRI signal correlates with both the contrast sensitivity and the macular thickness per quadrant. In contrast, in controls with simulated VFDs this correlation is absent. Moreover, our results support the occurrence of glaucomatous alterations beyond the scotomatic regions, apparent from decreased BOLD activity throughout the primary visual cortex. These functional alterations extend beyond V1 and affect also V2 and V3. Our findings may be explained by transsynaptic degeneration, in which damaged nerves in the optic nerve and the subsequent death of retinal ganglion cells will affect the entire visual pathway (including the optic nerve, lateral geniculate nucleus, optic radiation) and visual cortex, resulting in fairly widespread neuronal loss ^37–39^. Our findings of reduced BOLD modulation in glaucoma participants are in agreement with those of previous studies ^40–42^, even though one other study found no association between the BOLD signal and RNFL thickness ^43^. The reduced BOLD amplitude that we measured in early visual cortex may also be the result of deficient cortical perfusion, an explanation that would be in line with previous studies which showed that participants with glaucoma have reduced cerebral blood flow in early visual cortical areas when compared to controls ^44, 45^. The question remains whether limitations in perfusion could result in reduced functionality and evoke neural degeneration.

Based on a retinotopic mapping analysis, we found that overall, in the participants with glaucoma, their cortical areas retain their coarse retinotopic organization. Locally, however, their cortices have deviant neural configurations as evident from differences in sizes and positions of the pRFs. Importantly, these differences were present in comparison to control participants with matched simulated scotomas. But, the pattern of changes varied substantially between participants with glaucoma. Given that our glaucoma population has heterogenous VFD that are located at different positions of the VF and variable extent, a group level analysis would be insensitive to any local scotoma specific changes. This therefore called for a participant-specific analysis, in which the differences in pRF estimates within each glaucoma-control pair were compared to the expected intersubject variability in pRF properties observed in controls. This analysis, presented in detail in the SI, indicated that each glaucoma participant is unique in their local RF reconfigurations. While for some participants the reconfiguration is manifested by changes in the size of the pRFs, in others it manifests in different positions of the pRFs.

Overall, the size of pRFs and the deviations in their position distributions increases with larger damage to the VF (loss in contrast sensitivity as assessed via SAP). Notably, the same effect was not observed in the controls with simulated scotoma. We interpret these differences in pRFs properties as evidence for limited local cortical plasticity in adults with glaucoma. These local changes in pRF properties are consistent with what was previously described in homonymous VFDs ^15, 36^. This reorganization may involve the activation of long-range horizontal connections in the visual cortex, such that healthy neurons in the cortex surrounding the lesion thus take control of the deprived ones. These reconfigurations enable the neurons within the lesion projection zone to capture information from spared portions of the VF ^4, 11, 12, 14^. This not only holds true for the visual cortex, but also for other sensory areas of the adult cortex ^46–49^, cortical maps can for instance also reorganize after loss of sensory afferent nerves from a limb ^50^. Indeed, such capturing of information from outside the lesion projection zone would be required for predictive masking of the natural scotomas to occur, a phenomenon that is clinically frequently observed in glaucoma ^22^.

Furthermore, the VF reconstructions based on the fMRI data show that the cortical sensitivity of the glaucoma participants is reduced compared to the controls with simulated scotomas. This suggests that the glaucomatous deterioration of the visual system goes beyond the retina or that the way that the visual input is processed at the level of the retina is altered in glaucoma. In addition, while some glaucoma participants exhibit a lower VF sensitivity mapped via fMRI compared to what was measured with SAP (including damage in regions of the VF that appear to be 100% functional in SAP) others seem to show much smaller scotomas than what was measured via SAP. There are multiple explanatory mechanisms for these differences: 1) a reconfiguration of the RFs may cause neurons that were initially located inside the scotoma projection zone to process information from the spared VF, resulting in predictive masking of natural scotomas, and therefore the scotomas become smaller than measured in via SAP ^51^, 2) fMRI allows to detect subtle changes in the visual system functioning to which SAP is insensitive and 3) the accuracy of standard automated perimetry (SAP) measures was insufficient. Furthermore, the scotoma projection zone shrinks across the visual hierarchy. This finding supports the view that predictive masking of the scotoma (the reason why glaucoma participants cannot perceive their scotomas), may result from feedback from higher cortical areas where scotoma representation is small or inexistent to earlier visual areas. This mechanism can also be the driving force behind pRF shifts ^35^.

Our findings show that visual cortex functioning is altered in glaucoma. Glaucoma participants show reduced BOLD modulation compared controls; pRF size and position deviations are larger in quarterfield sections that showed greater loss in contrast sensitivity, and VF sensitivity of glaucoma participants differ from controls SS. These cerebral adaptations have important applications as the clinical diagnosis and treatments for glaucoma are currently only focused on the eye. The potential involvement of the brain and its plastic mechanisms in glaucoma suggests that the diagnosis should involve the assessment of neuronal function and the treatment should consider the entire visual pathway ^52–54^.

Evaluating the presence of neuroplasticity requires accurate and complex experimental conditions (e.g. using matched SS). While we attempted to match the visual input between participants with glaucoma and their matched control participants in the most accurate way possible, there are limitations associated with the simulation of the VFDs based on the contrast sensitivity measured using SAP. Errors in the assessment of the contrast sensitivity might lead to inaccurate simulations and biases that, in turn, may erroneously be interpreted as signs of reorganization. Nevertheless, we are convinced that this does not affect our present conclusions. In a previous study, we reduced the retinotopic stimulus contrast to 2% (from the conventional 50%). This reduced BOLD modulation did not affect the pRF estimates in early visual areas ^34^. While inaccurate SAP measurement may lead to somewhat inaccurate simulations, these are unlikely to introduce strong biases in the pRF estimates.

The participants with glaucoma were heterogeneous in various aspects, for instance in the extent of their scotoma, their disease duration, and in the asymmetry of the VFDs between both eyes. Such differences may affect the degree of neural reorganization as well as the degree of predictive masking taking place. Inside the scanner, only a relatively limited central part of the VF could be stimulated. Yet, in glaucoma, VFDs originate in the periphery of the visual field and many of the foveal scotomas that we could assess had relatively little reductions in contrast sensitivity. Nevertheless, we found marked differences in BOLD response, amongst others. Still, for this reason, future studies could consider including participants with more advanced glaucoma that would also have binocularly overlapping scotomas in their central vision. This could help to further establish the relation between the severity of the disease, the magnitude of the BOLD signal and any deviations in pRF properties. This information would also be beneficial when assessing the accuracy of the VF reconstructions. Alternatively, or additionally, it could be useful to perform the studies using visual stimulation that could reach deeper into the visual periphery ^55^.

The specific origin of the pRF reorganization patterns is not known nor are the factors that determine which specific adaptations take place (position shifts or size changes), this implies that some major points need to be considered. First, the techniques that we use reflect the aggregate RF properties at the population and/or subpopulation level. The pRF dynamics that we measure could result from changes in a subset of neurons ^17^. Second, changes in pRF properties may also result from extra-classical RF modulations and from attentional modulation. Such interactions can be studied by applying advanced neural computational models which have the ability to capture the activity of multiple subpopulations ^31, 34^ to take into account the extra classical RF representation ^56^; to model the effect of attention and higher order cognitive functions ^57^ and using stimuli that target specific neural populations ^31, 34^. Stimulus-driven approaches, such as pRF mapping, have an inherent disadvantage when applied to study neuroplasticity after VF loss. As we have seen in our present study, despite adding accurate control conditions, differences in the visual input of participants with glaucoma and control participants may still potentially influence results ^58^. One way to circumvent this limitation is to apply cortico-cortical models which are stimulus-agnostic by design. When applied to resting state data an approach such as connective field modeling may be used to evaluate if similar patterns of reorganization can be found in the absence of stimulation ^59, 60^.

Although this study sheds light on the neural mechanisms underlying predictive masking of natural scotomas and cortical reorganization, the entity and mechanism of cortical reorganization are still not completely clear. As considerable reorganization can be expressed subcortically, this should be studied using ultra high fields fMRI where different layers across the cortex can be measured. This will be important to quantify the level of adult brain plasticity in visual processing. Moreover, the VF reconstruction based on fMRI will improve using higher spatial resolution scans.

## 4. Conclusion

Participants with glaucoma exhibit individually unique patterns of reorganization that, nevertheless, suggest that the primary visual cortex of adults with glaucoma retain a local and limited degree of reorganization. This manifested itself in shifts in the centers of pRFs as well as in changes in their sizes. For some participants with glaucoma these changes even extend beyond their scotoma projection zone. Such changes to the neuronal configuration of the RFs may contribute to the masking of VFD which prevents patients from noticing their VFD. Moreover, although limited in spatial extent, this neuroplasticity may be critical to the successful implementation of future restorative therapies, e.g. those based on stem-cells.

## 5. Methods

### 5.1 Study Population

Nineteen individuals with primary open angle glaucoma (POAG) and nineteen control participants with normal or corrected vision were recruited. The population demographics are shown in table 1. The monocular data acquired for these participants was previously included in another study ^61^, so the details about the inclusion and exclusion criteria as well as the details of the ophthalmic data acquisition can also be found at ^61^. Here we focused on the acquired binocular data. Each age-matched control was assigned to a participant with glaucoma. This pairing was done based on demographic parameters such as age and gender. Prior to the ophthalmologic assessment, participants signed an informed consent form. Experimental protocols were approved by the University Medical Center Groningen, Medical Ethical Committee and conducted in accordance with the Declaration of Helsinki.

**Table 1.** Demographics of participants with glaucoma and controls. Average and standard deviation of age, percentage of female, intraocular pressure (IOP) for the right and left eye (average over three measurements), peripapillary retinal nerve fiber layer (pRNFL) thickness and the VF mean deviation (VFMD) measured with SAP for the right and left eye. Note that participants with glaucoma were receiving treatment.

| Measure | Participants with Glaucoma |  | Controls |  |
| --- | --- | --- | --- | --- |
|  | Average (n=19) | Standard Deviation (n=19) | Average (n=19) | Standard Deviation (n=19) |
| Age (y) | 70 | 8.8 | 68 | 7.3 |
| Gender (F%) | 52 | - | 42 | - |
| IOP (R/L; mmHg) | 13.4/13.4 | 2.6/3.9 | 13.1/13.5 | 2.8/3.2 |
| pRNFL thickness | 72.7/ 68.0 | 11.7/9.6 | 96.84/97.2 | 9.7/10.9 |

**Table 1. Demographics of participants with glaucoma and controls.**
| (R/L; $\mu\text{m}$) | | | | |
| --- | --- | --- | --- | --- |
| VFMD (R/L; dB) | -7.4/-9.1 | 8.1/8.3 | -0.4/-0.69 | 1.4/3.4 |

Inclusion criteria for the participants with glaucoma were as follows: having an intraocular pressure (IOP) > 21 mmHg before treatment onset, the presence of a VFD due to glaucoma (glaucoma hemifield test outside normal limits), abnormal optical coherence tomography (OCT); peripapillary retinal nerve fiber layer thickness (pRNFL) at least one clock hour with a probability <0.01, spherical equivalent refraction within ±3 D.

Exclusion criteria for both groups were: having any ophthalmic disorder affecting visual acuity or VF (other than primary open angle glaucoma (POAG) in the participants with glaucoma), any neurological or psychiatric disorders, the presence of gross abnormalities or lesions on their magnetic resonance imaging (MRI) scans, or having any contraindication for MRI (e.g., having a pacemaker or being claustrophobic).

### 5.2 Ophthalmic data

Prior to their participation in the MRI experiments, we assessed for all participants their visual acuity, IOP, VF sensitivity (measured using HFA and frequency doubling technology [FDT]) and retinal nerve fiber layer (RNFL) thickness. Visual acuity was measured using a Snellen chart with optimal correction provided for the viewing distance. IOP was measured using a Tonoref noncontact tonometer (Nidek, Hiroishi, Japan). The VFs were first screened using FDT (Carl Zeiss Meditec) using the C20-1 screening mode. The contrast sensitivity at several locations of the VF was measured using SAP specifically using a HFA (Carl Zeiss Meditec, Jena, Germany) with the 24-2 or 30-2 grid and the Swedish Interactive Threshold Algorithm (SITA) Fast. Only reliable HFA tests were included in this study. A VF test result was considered unreliable if false-positive errors exceeded 10% or fixation losses exceeded 20% and false-negative errors exceeded 10% ^62^. Finally, the RNFL thickness was measured by means of OCT using a Canon OCT-HS100 scanner (Canon, Tokyo, Japan).

### 5.3 Experimental Procedure

Each participant completed two (f)MRI sessions of approximately 1.5h each. In the first session, the anatomical scan (T1w), Diffusion Weighted Imaging (DWI), T2w, resting state functional scans and a MT localizer were acquired. In the second session, the retinotopic mapping and scotoma localizers experiments took place. These experiments were performed binocularly and monocularly as well. The resting state fMRI results are reported on in a different paper ^63^. Here, we report on the results of the binocular retinotopic mapping scans.

#### 5.3.1 Participants with glaucoma

The second session differed for participants with glaucoma and control participants. For the participants with glaucoma, their second (f)MRI session comprised the retinotopy and scotoma localizer experiments. The retinotopy experiment comprised nine runs in total of which six were done with binocular and three with monocular vision. For this study, only the binocular runs were analyzed. The scotoma localizer experiment comprised 2 runs in total of which one was done with binocular and one with monocular vision. This task was performed to control residual activity within the scotoma projection zone. In the monocular experiments, the most lesioned eye was stimulated and the other was occluded using an MRI compatible opaque lense. The most lesioned eye was selected based on the SAP MD (mean deviation) score; the eye with the lowest MD was selected. The monocular retinotopy results were used to assess the capability of fMRI to detect visual field defects and are reported on in a different paper ^61^.

#### 5.3.2 Control participants

In their second (f)MRI session, the control participants performed the LCR, LCR SS and scotoma localizer experiments. The latter was used to define the simulated scotoma projection zone. All experiments were done with binocular vision. For both LCR and LCR SS, four runs were performed.

Two scotoma localizers were acquired, one with and another without the SS superimposed on the stimulus.

### 5.4 Stimulus presentation and image acquisition

Stimuli were presented on an MR compatible display screen (BOLDscreen 24 LCD; Cambridge Research Systems, Cambridge, UK). The screen was located at the head-end of the MRI scanner. Participants viewed the screen through a tilted mirror attached to the head coil. Distance from the participant’s eyes to the display (measured through the mirror) was 120 cm. Screen size was 22x14 deg. The maximum stimulus radius was 7 deg of visual angle. Visual stimuli were created using MATLAB (Mathworks, Natick, MA, USA) and the Psychtoolbox ^64, 65^.

#### 5.4.1 Stimuli

All participants underwent binocular visual field mapping using luminance contrast retinotopy (LCR) mapping. Figure 8A shows an example frame of the stimulus. Additionally, the glaucoma participants observed the LCR monocularly, and the healthy participants viewed the LCR binocularly with a simulated scotoma (LCR SS) superimposed (Figure 8B). For each control, the LCR SS was matched to that of a participant with glaucoma (see section 5.4.1.2). This condition acted as a reference for the glaucoma binocular LCR, in order to disentangle possible (cortical) plasticity.

**Figure 8.**
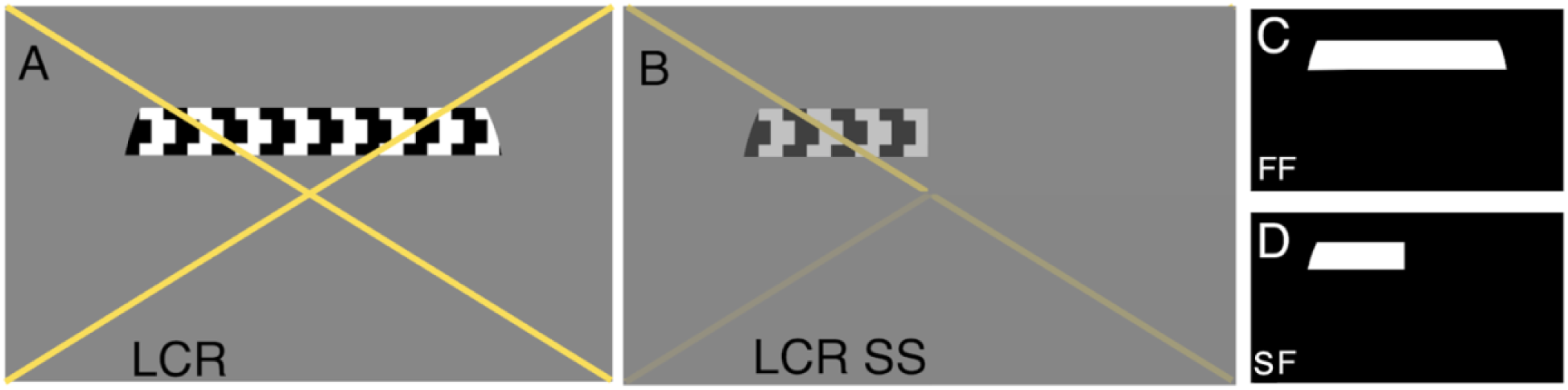
Example of the stimuli used to obtain pRF parameter estimates. (A) LCR stimulus. **(B)** LCR SS stimulus, this particular example depicts the contrast sensitivity loss of participant G01 (MD(OS)=-14.72 MD (OD)=-8.37). The colour of the cross changed between yellow and black, and served to guide the gaze of the participant with central scotomas to the center of the cross. Panels C and D show the full field (FF) the scotoma field (SF) models used in the pRF analysis, respectively.

##### 5.4.1.1 Luminance-contrast retinotopy (LCR)

LCR consisted of a drifting bar aperture defined by high-contrast flickering texture ^66^. The bar aperture, i.e. alternating rows of high-contrast luminance checks drifting in opposite directions, moved in eight different directions: four bar orientations (horizontal, vertical, and the two diagonal orientations) and for each orientation two opposite drift directions. The bar moved across the screen in 16 equally spaced steps, each lasting 1 TR (repetition time, time between two MRI volume acquisitions). The bar contrast, width, and spatial frequency were 100%, 1.75 degree, and 0.5 cycles per degree, respectively. After each pass, during which the bar moved across the entire screen during 24 s, the bar moved across half of the screen for 12 s, followed by a blank full screen stimulus at mean luminance for 12 s as well.

##### 5.4.1.2 Luminance-contrast defined retinotopy with simulated scotomas (LCR SS)

LCR SS consisted of the LCR stimulus with a simulated scotoma. The SS for a control participant was designed to mimic the contrast sensitivity of the corresponding glaucoma participant under binocular vision. The scotoma was simulated by means of local reductions in stimulus contrast. In particular, the SS consisted of an alpha transparency contrast layer defined using the HFA sensitivity values of the respective participant with glaucoma. For example, a decrease of 3dB in HFA sensitivity is simulated by means of a reduction in stimulus contrast of 50%. The binocular HFA sensitivity at every position measure was calculated by taking the maximum between Left and Right eye.

##### 5.4.1.3 Attentional task

During the retinotopic mapping scans, participants were required to perform a fixation task in which they had to press a button each time the fixation cross changed colour between black and yellow (retinotopic experiments) and between white and black (scotoma localizer). The fixation cross extended towards the edges of the screen so that it could be used as a queue for the screen’s center by the participants with central scotomas. The average performance was above 75% for all conditions and for participants with glaucoma and control participants. The task performance per condition is presented in table S1.

#### 5.4.2 Magnetic resonance imaging

##### 5.4.2.1 Data acquisition and preprocessing

Scanning was carried out on a 3 Tesla Siemens Prisma MR-scanner using an 64-channel receiving head coil. A T1-weighted scan (voxel size, 1mm^3^; matrix size, 256 x 256 x 256) covering the whole-brain was recorded to chart each participant’s cortical anatomy. Padding was applied to strike a balance between participant comfort and a reduction of possible head motion. The retinotopic scans were collected using standard EPI sequence (TR, 1500 ms; TE, 30 ms; voxel size, 3mm^3^, flip angle 80; matrix size, 84 x 84 x 24). Slices were oriented to be approximately parallel to the calcarine sulcus. For all retinotopic scans (LCR, LCR monocular and LCR SS), a single run consisted of 136 functional images (duration of 204 s). The (S)SPZ localizers consisted of 144 functional images (duration of 216 s).

The T1-weighted whole-brain anatomical images were reoriented in AC-PC space. The resulting anatomical image was initially automatically segmented using Freesurfer ^67^ and subsequently edited manually. The cortical surface was reconstructed at the gray/white matter boundary and rendered as a smoothed 3D mesh ^68^.

The functional scans were analysed in the mrVista software package for MATLAB (available at https://web.stanford.edu/group/vista/cgi-bin/wiki/index.php/MrVista). Head movement between and within functional scans were corrected ^69^. The functional scans were averaged and coregistered to the anatomical scan ^69^, and interpolated to a 1mm isotropic resolution. Drift correction was performed by detrending the BOLD time series with a discrete cosine transform filter with a cutoff frequency of 0.001Hz. In order to avoid possible saturation effects, the first 8 images were discarded.

##### 5.4.2.2 Visual field mapping and ROI definition

The pRF analysis was performed using both conventional population receptive field (pRF) mapping^66^ and micro-probing ^70^. Using both models, for all the participants the functional responses to binocular LCR were analysed using a full field (FF) model (Figure 8C). Additionally, the data acquired in LCR SS condition in the control participants in experiments 1 and 2 were analyzed using a model that included the simulated scotoma (scotoma field; SF, Figure 8D).

The visual areas V1, V2 and V3 were defined on the basis of phase reversal on the inflated cortical surface, obtained with the conventional pRF model using the LCR stimulus presented binocularly.

### 5.5 Population receptive field analysis

As in previous work ^15, 66, 71^ data was thresholded by retaining the pRF models that explained at least 0.15 of the variance in the BOLD response and that had an eccentricity in the range of 0-7 degrees, for all conditions (i.e., LCR and LCR SS).

#### 5.5.1 Correlation Analysis

The correlations between the BOLD modulation, the pRF size and position deviation and the disease severity were calculated using a linear mixed effects model with a slope and intercept per subject as a random effect.

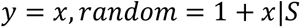

Where *y* is the dependent variable, i.e BOLD modulation, pRF size and position deviation, *x* is the independent variable, i.e contrast sensitivity and MNFL, and S is the subject. To determine whether the BOLD modulation, defined as the standard deviation of the BOLD signal, of glaucoma participants was different from control participants, repeated measures two-way analysis of variance (ANOVA) with ROIs, eccentricity and condition (LCR and LCR SS) were performed.

#### 5.5.2 Analysis of the variation in pRF properties between glaucoma and control participants

To investigate changes in pRF sizes between glaucoma and control participants the cumulative distribution of pRF size is depicted across voxels. At the group level, statistical testing was performed by calculating the median pRF size across voxels per participant and comparing these median values across groups (glaucoma vs control participants) using a two sample t-test.

Given the heterogeneity of the VFD of the glaucoma participants, one of the challenges is how to properly perform a group level analysis. Analyzing the pRF deviation at the level of quarter fields and correlating these deviations with SAP and OCT metrics, allows us to understand if at the group level, the deviations in pRF positions are related to disease severity. In order to understand how pRF properties are associated with the severity of glaucoma, we performed two different analysis: 1) we correlated the averaged pRF size and ED of the average position calculated per quarter field between participants with glaucoma and respective control participant with the contrast sensitivity and MNFL thickness; and 2) we assessed the significance of the deviation between pairs in relation to an expected baseline deviation, which summarizes the variation in pRF properties amongst control participants. In order to take this into account, a group level analysis was done by determining the rank of the deviation of the participant pair (either the glaucoma-control SS pair or the control SS-control NS pair) relative to the baseline deviation (which consisted of all the 19 deviations between the matching control participant and all other control participants, Figure S10). This approach and its results are presented in detail in the sections 7 and 8 of the supplementary information.

#### 5.5.3 Visual field reconstruction analysis

Both MP and pRF mapping techniques allow an accurate reconstruction of the VF by back-projecting the pRF properties of all voxels within a visual area onto the VF ^61, 72^. The VF reconstructed maps reflect the VF sampling density. Since the presence of VFD reduces the sampling of a particular region of the VF, fMRI-based VF reconstruction techniques are suitable to detect VFDs and indirectly reflect VF sensitivity.

In this study we compared the visual field reconstruction between participants with Glaucoma and Controls SS. The visual field reconstruction was obtained using the MP technique as described in ^61^. In addition, to directly compare the VF reconstructions between groups, VF maps were converted from normalized scale to a dB scale by taking the 10 × log10 of the sampling density values, resulting in VF sensitivity values. The final VF maps correspond to the deviation of VF sensitivities between glaucoma and controls SS from controls NS according to the 90, 95, 98, 99 and 99.5 % CI boundaries. This approach was previously applied by ^72^.

#### 5.5.4 Statistical Analysis

All statistical analyses were performed using R (version 0.99.903; R Foundation for Statistical Computing, Vienna, Austria) and MATLAB (version 2016b; Mathworks, Natick, MA, USA). After correction for multiple comparisons, a p-value of 0.05 or less was considered statistically significant.

## Acknowledgments

We would like to thank Prof. Nomdo Jansonius for providing access to participants with glaucoma and his feedback regarding how to simulate the visual field defects experienced by the glaucoma participants in case-matched controls. We would also like to thank Hinke Halbertsma for her help and fruitful discussions regarding the visualization of the visual field backprojections. Authors JC and AI were supported by the European Union’s Horizon 2020 research and innovation programme under the Marie Sklodowska-Curie grant agreements No. 641805 (NextGenVis) and No. 661883 (EGRET). AI, and JC received additional funding from the Graduate School of Medical Sciences (GSMS), University of Groningen, The Netherlands. The funding organization had no role in the design, conduct, analysis, or publication of this research.

## Author contributions

JC and FWC conceptualized and designed the study. JC and AI performed the data collection. JC and JM performed the data analysis. RJR provided feedback on the methodology. FC, NMJ and RR supervised the work. JC wrote the first draft manuscript. All authors contributed to manuscript revision, read, and approved the submitted version.

## Data availability statement

The raw data supporting the conclusions of this article will be made available by the authors.

The code for the Micro Probing based mapping of RFs and visual field reconstruction is available via the following links: https://github.com/Joana-Carvalho/Micro-Probing; https://github.com/Joana-Carvalho/FMRI-based-Visual-field-Reconstruction

## References

1. Haak, K. V. et al. Abnormal visual field maps in human cortex: a mini-review and a case report. Cortex 56, 14–25 (2014).

2. Halbertsma, H. N., Haak, K. V. & Cornelissen, F. W. Stimulus- and Neural-Referred Visual Receptive Field Properties following Hemispherectomy: A Case Study Revisited. Neural Plast. 2019, 6067871 (2019).

3. Hoffmann, M. B. & Dumoulin, S. O. Congenital visual pathway abnormalities: a window onto cortical stability and plasticity. Trends Neurosci. 38, 55–65 (2015).

4. Dilks, D. D., Baker, C. I., Peli, E. & Kanwisher, N. Reorganization of Visual Processing in Macular Degeneration Is Not Specific to the ‘Preferred Retinal Locus’. J. Neurosci. 29, 2768–2773 (2009).

5. Clavagnier, S., Dumoulin, S. O. & Hess, R. F. Is the Cortical Deficit in Amblyopia Due to Reduced Cortical Magnification, Loss of Neural Resolution, or Neural Disorganization? J. Neurosci. 35, 14740– 14755 (2015).

6. Hoffmann, M. B., Tolhurst, D. J., Moore, A. T. & Morland, A. B. Organization of the visual cortex in human albinism. J. Neurosci. 23, 8921–8930 (2003).

7. Morland, A. B. & Hoffmann, M. Retinotopic organisation of the visual cortex in human albinism. Journal of Vision vol. 2 46–46 (2002).

8. Muckli, L., Naumer, M. J. & Singer, W. Bilateral visual field maps in a patient with only one hemisphere. Proceedings of the National Academy of Sciences vol. 106 13034–13039 (2009).

9. Levin, N., Dumoulin, S. O., Winawer, J., Dougherty, R. F. & Wandell, B. A. Cortical maps and white matter tracts following long period of visual deprivation and retinal image restoration. Neuron 65, 21– 31 (2010).

10. Baseler, H. A. et al. Reorganization of human cortical maps caused by inherited photoreceptor abnormalities. Nat. Neurosci. 5, 364–370 (2002).

11. Gilbert, C. D. & Wiesel, T. N. Receptive field dynamics in adult primary visual cortex. Nature 356, 150–152 (1992).

12. Schumacher, E. H. et al. Reorganization of visual processing is related to eccentric viewing in patients with macular degeneration. Restor. Neurol. Neurosci. 26, 391–402 (2008).

13. Dilks, D. D., Baker, C. I., Peli, E. & Kanwisher, N. Reorganization of visual processing in macular degeneration is not specific to the ‘preferred retinal locus’. Journal of Vision vol. 9 732–732 (2010).

14. Pettet, M. W. & Gilbert, C. D. Dynamic changes in receptive-field size in cat primary visual cortex. Proc. Natl. Acad. Sci. U. S. A. 89, 8366–8370 (1992).

15. Baseler, H. A. et al. Large-scale remapping of visual cortex is absent in adult humans with macular degeneration. Nat. Neurosci. 14, 649–655 (2011).

16. Borges, V. M., Danesh-Meyer, H. V., Black, J. M. & Thompson, B. Functional effects of unilateral open-angle glaucoma on the primary and extrastriate visual cortex. J. Vis. 15, 9 (2015).

17. Haak, K. V., Cornelissen, F. W. & Morland, A. B. Population receptive field dynamics in human visual cortex. PLoS One 7, e37686 (2012).

18. Prabhakaran, G. T. et al. Foveal pRF properties in the visual cortex depend on the extent of stimulated visual field. Neuroimage 222, 117250 (2020).

19. Kingman, S. Glaucoma is second leading cause of blindness globally. Bull. World Health Organ. 82, 887–888 (2004).

20. Crabb, D. P., Smith, N. D., Glen, F. C., Burton, R. & Garway-Heath, D. F. How Does Glaucoma Look? Ophthalmology vol. 120 1120–1126 (2013).

21. Hu, C. X. et al. What do patients with glaucoma see? Visual symptoms reported by patients with glaucoma. Am. J. Med. Sci. 348, 403–409 (2014).

22. Hoste, A. M. New insights into the subjective perception of visual field defects. Bull. Soc. Belge Ophtalmol. 65–71 (2003).

23. Duncan, R. O., Sample, P. A., Weinreb, R. N., Bowd, C. & Zangwill, L. M. Retinotopic organization of primary visual cortex in glaucoma: Comparing fMRI measurements of cortical function with visual field loss. Prog. Retin. Eye Res. 26, 38–56 (2007).

24. Duncan, R. O., Sample, P. A., Weinreb, R. N., Bowd, C. & Zangwill, L. M. Retinotopic organization of primary visual cortex in glaucoma: a method for comparing cortical function with damage to the optic disk. Invest. Ophthalmol. Vis. Sci. 48, 733–744 (2007).

25. Zhou, W. et al. Retinotopic fMRI Reveals Visual Dysfunction and Functional Reorganization in the Visual Cortex of Mild to Moderate Glaucoma Patients. J. Glaucoma 26, 430–437 (2017).

26. Wandell, B. A. & Smirnakis, S. M. Plasticity and stability of visual field maps in adult primary visual cortex. Nat. Rev. Neurosci. 10, 873–884 (2009).

27. Calford, M. B. et al. Neuroscience: rewiring the adult brain. Nature vol. 438 E3; discussion E3–4 (2005).

28. Smirnakis, S. M. et al. Lack of long-term cortical reorganization after macaque retinal lesions. Nature 435, 300–307 (2005).

29. Binda, P., Thomas, J. M., Boynton, G. M. & Fine, I. Minimizing biases in estimating the reorganization of human visual areas with BOLD retinotopic mapping. J. Vis. 13, 13 (2013).

30. Hoffmann, M. B., Schmidtborn, L. C. & Morland, A. B. [Abnormal representations in the visual cortex of patients with albinism: diagnostic aid and model for the investigation of the self-organisation of the visual cortex]. Ophthalmologe 104, 666–673 (2007).

31. Carvalho, J. et al. Micro-probing enables fine-grained mapping of neuronal populations using fMRI. Neuroimage 209, 116423 (2020).

32. Redmond, T., Garway-Heath, D. F., Zlatkova, M. B. & Anderson, R. S. Sensitivity loss in early glaucoma can be mapped to an enlargement of the area of complete spatial summation. Invest. Ophthalmol. Vis. Sci. 51, 6540–6548 (2010).

33. Rountree, L. et al. Optimising the glaucoma signal/noise ratio by mapping changes in spatial summation with area-modulated perimetric stimuli. Sci. Rep. 8, 2172 (2018).

34. Yildirim, F., Carvalho, J. & Cornelissen, F. W. A second-order orientation-contrast stimulus for population-receptive-field-based retinotopic mapping. Neuroimage 164, 183–193 (2018).

35. Carvalho, J., Renken, R. J. & Cornelissen, F. W. Predictive masking of an artificial scotoma is associated with a system-wide reconfiguration of neural populations in the human visual cortex. NeuroImage vol. 245 118690 (2021).

36. Papanikolaou, A. et al. Population receptive field analysis of the primary visual cortex complements perimetry in patients with homonymous visual field defects. Proc. Natl. Acad. Sci. U. S. A. 111, E1656–65 (2014).

37. Prins, D., Hanekamp, S. & Cornelissen, F. W. Structural brain MRI studies in eye diseases: are they clinically relevant? A review of current findings. Acta Ophthalmol. 94, 113–121 (2016).

38. Gupta, N., Ang, L.-C., de Tilly, L. N., Bidaisee, L. & Yücel, Y. H. Human glaucoma and neural degeneration in intracranial optic nerve, lateral geniculate nucleus, and visual cortex. Br. J. Ophthalmol. 90, 674–678 (2006).

39. Haykal, S., Curcic-Blake, B., Jansonius, N. M. & Cornelissen, F. W. Fixel-Based Analysis of Visual Pathway White Matter in Primary Open-Angle Glaucoma. Invest. Ophthalmol. Vis. Sci. 60, 3803–3812 (2019).

40. Wuerger, S., Powell, J., Chaudhoury, A. & Parkes, L. Comparison of fMRI measurements in LGN and Primary Visual cortex with visual deficits in Glaucoma. Journal of Vision vol. 15 257 (2015).

41. Duncan, R. O., Sample, P. A., Bowd, C., Weinreb, R. N. & Zangwill, L. M. Arterial spin labeling fMRI measurements of decreased blood flow in primary visual cortex correlates with decreased visual function in human glaucoma. Vision Res. 60, 51–60 (2012).

42. Murphy, M. C. et al. Retinal Structures and Visual Cortex Activity are Impaired Prior to Clinical Vision Loss in Glaucoma. Sci. Rep. 6, 31464 (2016).

43. Qing, G., Zhang, S., Wang, B. & Wang, N. Functional MRI signal changes in primary visual cortex corresponding to the central normal visual field of patients with primary open-angle glaucoma. Invest. Ophthalmol. Vis. Sci. 51, 4627–4634 (2010).

44. Wang, Q. et al. Reduced Cerebral Blood Flow in the Visual Cortex and Its Correlation With Glaucomatous Structural Damage to the Retina in Patients With Mild to Moderate Primary Open-angle Glaucoma. Journal of Glaucoma vol. 27 816–822 (2018).

45. Zhang, S. et al. Retinotopic Changes in the Gray Matter Volume and Cerebral Blood Flow in the Primary Visual Cortex of Patients With Primary Open-Angle Glaucoma. Invest. Ophthalmol. Vis. Sci. 56, 6171–6178 (2015).

46. Kaas, J. H. Sensory loss and cortical reorganization in mature primates. Prog. Brain Res. 138, 167–176 (2002).

47. Jain, N., Qi, H.-X., Collins, C. E. & Kaas, J. H. Large-scale reorganization in the somatosensory cortex and thalamus after sensory loss in macaque monkeys. J. Neurosci. 28, 11042–11060 (2008).

48. Mühlnickel, W., Elbert, T., Taub, E. & Flor, H. Reorganization of auditory cortex in tinnitus. Proc. Natl. Acad. Sci. U. S. A. 95, 10340–10343 (1998).

49. Kambi, N. et al. Large-scale reorganization of the somatosensory cortex following spinal cord injuries is due to brainstem plasticity. Nat. Commun. 5, 3602 (2014).

50. Makin, T. R. & Flor, H. Brain (re)organisation following amputation: Implications for phantom limb pain. Neuroimage 218, 116943 (2020).

51. Carvalho, J., Renken, R. J. & Cornelissen, F. W. Predictive masking is associated with a system-wide reconfiguration of neural populations in the human visual cortex. doi:10.1101/758094.

52. Doozandeh, A. & Yazdani, S. Neuroprotection in glaucoma. Journal of Ophthalmic and Vision Research vol. 11 209 (2016).

53. Rusciano, D. et al. Neuroprotection in Glaucoma: Old and New Promising Treatments. Adv. Pharmacol. Sci. 2017, 4320408 (2017).

54. Nucci, C. et al. Neuroprotective agents in the management of glaucoma. Eye 32, 938–945 (2018).

55. Greco, V. et al. A low-cost and versatile system for projecting wide-field visual stimuli within fMRI scanners. Behav. Res. Methods 48, 614–620 (2016).

56. Zuiderbaan, W., Harvey, B. M. & Dumoulin, S. O. Modeling center-surround configurations in population receptive fields using fMRI. J. Vis. 12, 10 (2012).

57. Klein, B. P., Harvey, B. M. & Dumoulin, S. O. Attraction of position preference by spatial attention throughout human visual cortex. Neuron 84, 227–237 (2014).

58. Carvalho, J., Renken, R. J. & Cornelissen, F. W. Studying Cortical Plasticity in Ophthalmic and Neurological Disorders: From Stimulus-Driven to Cortical Circuitry Modeling Approaches. Neural Plast. 2019, 2724101 (2019).

59. Haak, K. V., et al. Connective field modeling. Neuroimage 66, 376–384 (2013).

60. Gravel, N. et al. Cortical connective field estimates from resting state fMRI activity. Front. Neurosci. 8, 339 (2014).

61. Carvalho, J. et al. Visual Field Reconstruction Using fMRI-Based Techniques. Transl. Vis. Sci. Technol. 10, 25 (2021).

62. Wesselink, C. & Jansonius, N. M. Glaucoma progression detection with frequency doubling technology (FDT) compared to standard automated perimetry (SAP) in the Groningen Longitudinal Glaucoma Study. Ophthalmic Physiol. Opt. 37, 594–601 (2017).

63. Demaria, G. et al. Binocular Integrated Visual Field Deficits Are Associated With Changes in Local Network Function in Primary Open-Angle Glaucoma: A Resting-State fMRI Study. Front. Aging Neurosci. 13, 744139 (2021).

64. Brainard, D. H. The Psychophysics Toolbox. Spat. Vis. 10, 433–436 (1997).

65. Pelli, D. G. The VideoToolbox software for visual psychophysics: transforming numbers into movies. Spat. Vis. 10, 437–442 (1997).

66. Dumoulin, S. O. & Wandell, B. A. Population receptive field estimates in human visual cortex. Neuroimage 39, 647–660 (2008).

67. Dale, A. M., Fischl, B. & Sereno, M. I. Cortical surface-based analysis. I. Segmentation and surface reconstruction. Neuroimage 9, 179–194 (1999).

68. Wandell, B. A., Chial, S. & Backus, B. T. Visualization and measurement of the cortical surface. J. Cogn. Neurosci. 12, 739–752 (2000).

69. Nestares, O. & Heeger, D. J. Robust multiresolution alignment of MRI brain volumes. Magn. Reson. Med. 43, 705–715 (2000).

70. Carvalho, J. et al. Micro-probing enables high-resolution mapping of neuronal subpopulations using fMRI. doi:10.1101/709006.

71. Winawer, J., Horiguchi, H., Sayres, R. A., Amano, K. & Wandell, B. A. Mapping hV4 and ventral occipital cortex: the venous eclipse. J. Vis. 10, 1 (2010).

72. Halbertsma, H. N., Bridge, H., Carvalho, J., Cornelissen, F. W. & Ajina, S. Visual Field Reconstruction in Hemianopia Using fMRI Based Mapping Techniques. Front. Hum. Neurosci. 15, 713114 (2021).

